# Sleep regulates the glial engulfment receptor Draper to promote Wallerian degeneration

**DOI:** 10.1101/716894

**Authors:** Bethany A. Stahl, James B. Jaggard, Alex C. Keene

## Abstract

Sleep, a universal behavior, is critical for diverse aspects of brain function. Chronic sleep disturbance is associated with numerous health consequences, including neurodegenerative disease and cognitive decline. Neurite damage due to apoptosis, trauma, or genetic factors is a common feature of aging, and clearance of damaged neurons is essential for maintenance of brain function. In the central nervous system, damaged neurites are cleared by Wallerian degeneration, in which activated microglia and macrophages engulf damaged neurons. The fruit fly *Drosophila melanogaster* provides a powerful model for investigating the relationship between sleep and Wallerian degeneration. Several lines of evidence suggest that glia influence sleep duration, sleep-mediated neuronal homeostasis, and clearance of toxic substances during sleep, raising the possibility that glial engulfment of damaged axons is regulated by sleep. To explore this possibility, we axotomized olfactory receptor neurons and measured the effects of sleep loss or gain on the clearance of damaged neurites. Mechanical sleep deprivation impaired the clearance of damaged neurites, whereas the sleep-promoting drug gaboxadol accelerated clearance. In sleep-deprived animals, multiple markers of glial activation were delayed, including activation of the JAK/STAT pathway, upregulation of the cell corpse engulfment receptor Draper, and innervation of the antennal lobe by glial membranes. These markers were all enhanced when sleep was induced in gaboxadol-treated flies. Taken together, these findings reveal a critical role for sleep in regulation glial activation and engulfment following axotomy, providing a platform for further investigations of the molecular mechanisms underlying sleep-dependent modulation of glial function and neurite clearance.

**Highlights:**

- Sleep deprivation impairs Wallerian degeneration in fruit flies.
- Pharmacological induction of sleep accelerates Wallerian degeneration.
- Sleep promotes innervation surrounding damaged neurites by phagocytic glia.
- Sleep increases levels of the glial activation markers Draper and Stat92E.

## Results and Discussion

Glial regulation of synaptic function and neuronal homeostasis is critical for proper sleep [1–3]. The *Drosophila* brain contains five distinct types of glia: perineural and subperineural glia, which form the blood-brain barrier; cortex glia; astrocyte-like glia, which regulate synaptic function; and ensheathing glia, which surround neuropil [4–6]. Multiple glial subtypes have been identified that regulate sleep in *Drosophila* [1–3]. Moreover, glial dysfunction is associated with aging, and numerous models of neurodegeneration in flies [7–11]. Despite these bidirectional interactions between glia and sleep, surprisingly little is known about the mechanisms through which sleep-deprivation impacts glial function, and how this contributes to neurodegenerative disease.

Experimental ablation of the antennae results in axotomy of the primary olfactory neurons, followed by engulfment of the damaged olfactory neurites by ensheathing glia [12]. This system allows precise quantification of glial engulfment, and has been used to identify novel genetic factors required for this process [13]. To investigate the relationship between Wallerian degeneration and sleep, we deprived flies of sleep by mechanical shaking following antennal axotomy (Fig 1A). For these experiments, we used a strain in which mcd8:GFP is expressed in Or22a neurons, thus labeling olfactory neurons [14]. Mechanical sleep deprivation resulted in a near complete loss of sleep during day and night, whereas antennal ablation had no effect on sleep in undisturbed flies (Fig S1A–C). Three days following axotomy, the remaining neurites in the antennal lobe of sleep-deprived flies or undisturbed controls were quantified. The signals from Or22a-labeled neurons were quantified independently in the axonal projections, antennal lobe, and antennal commissure (Fig S1D). Consistent with previous reports, axotomy resulted in a near complete loss of axons, as well as a significant reduction in olfactory receptor neuron projections with the antennal lobe (Fig 1B,C) [12,15]. Sleep deprivation did not affect axonal or glomerular innervation in intact flies, suggesting that sleep deprivation on its own does not impact olfactory neuron morphology (Fig 1D and Movie S1). Conversely, the signal from damaged neurites was significantly greater in sleep-deprived axotomized flies than in undisturbed controls (Fig 1E). Quantifications (Fig 1F–H) confirmed that fluorescence signals within the antennal glomeruli, antennal commissure, and neurites were all significantly greater in axotomized flies following sleep deprivation than in undisturbed axotomized controls.

**Figure 1.**
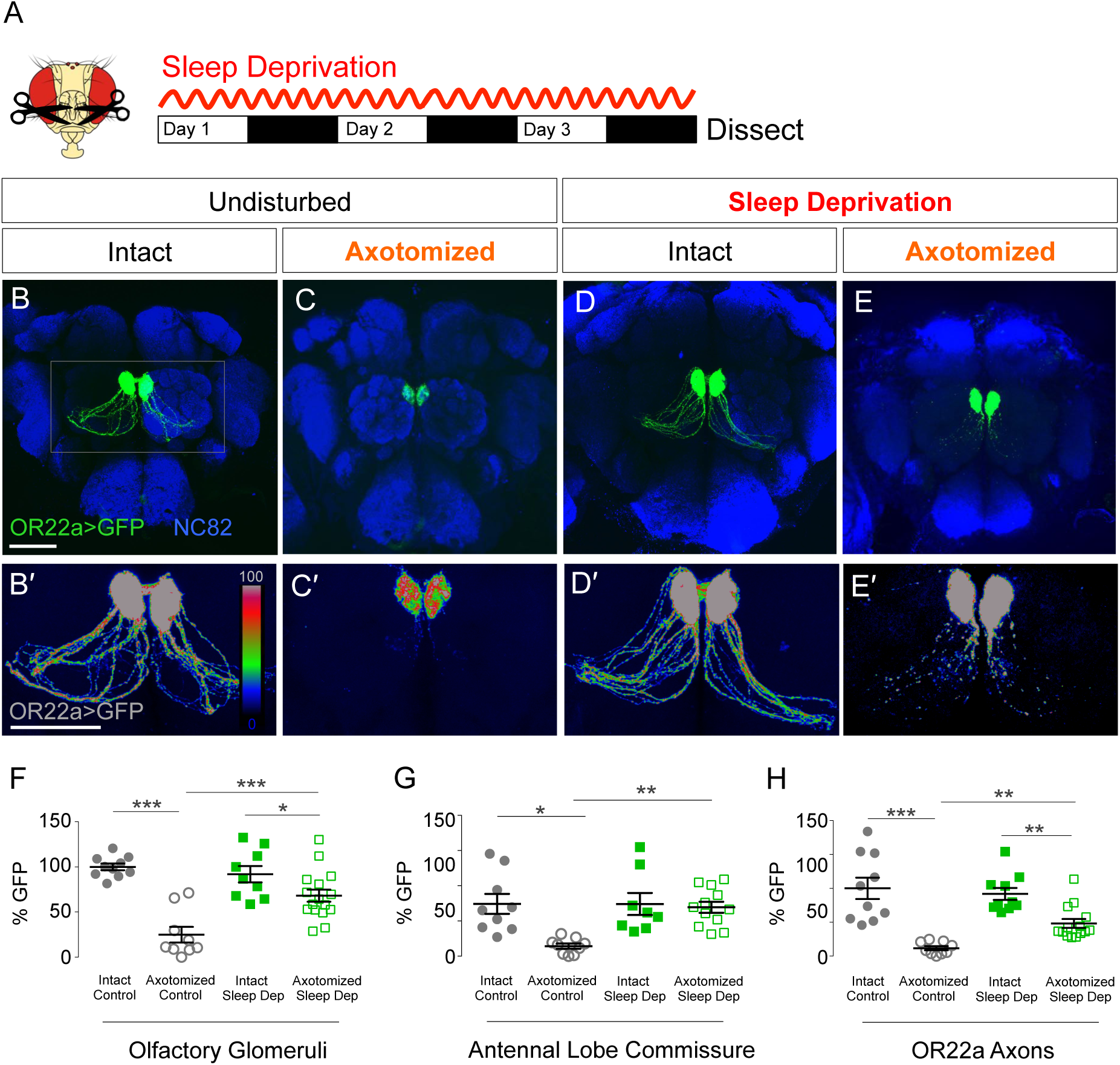
Sleep deprivation slows Wallerian degeneration. **(A)** Schematic of antennal ablation and sleep deprivation paradigm. **(B–E)** Maximum intensity projection of GFP-labeled olfactory OR22a axons (green) with NC82 neuropil counterstain (blue). **(B)** OR22a neurons in intact, undisturbed flies. **(C)** Flies with third antennal segment axotomoy and undisturbed treatment. **(D)** Sleep-deprived unharmed flies. **(E)** Flies subjected to ablation plus sleep deprivation treatment. **(B′–E′)** Zoom and max intensity rainbow plots. **(F–H)** In axotomized sleep-deprived flies, GFP signal was lower than in unharmed individuals, but clearance was slower than in axotomized controls in OR22a glomeruli (**F**; ANOVA, F_(3,40)_=18.03, P<0.0001), axons (**G**; ANOVA, F_(3,40)_=7.080, P<0.001) and the antennal lobe commissure (**H**; ANOVA, F_(3,40)_=16.72, P<0.0001). Tukey’s multiple comparisons tests: *P<0.05, **P<0.01, ***P<0.001. Scale Bar = 50 μm.

To validate these findings, we monitored GFP labeling from Or22a neurites over a time course following axotomy. In undisturbed flies, the signal was significantly reduced 1 day after axotomy and nearly absent after 4 days (Fig S1E–N). At all three time points (Day 2, Day 3, and Day 4), the GFP signal was greater in sleep-deprived axotomized flies than in undisturbed axotomized flies (Fig S1H–R). Together, these findings suggest that sleep deprivation impairs glial engulfment of axotomized neurons.

To determine whether sleep is sufficient to promote Wallerian degeneration, we pharmacologically induced sleep in axotomized flies and measured neurite clearance. In *Drosophila*, the GABA_A_ receptor agonist gaboxadol potently induces sleep ([16–18]; Fig S2A-C). Flies expressing cd8:GFP in Or22a neurons were placed on gaboxadol-laced food immediately following axotomy, and signal from the remaining neurites was measured on each of the four subsequent days. Gaboxadol treatment robustly increased daytime sleep in both intact and axotomized flies, with no difference in total sleep duration between the two groups (Fig S2A–C). Gaboxadol treatment had no effect on the GFP signal from Or22a neurons in intact flies (Fig 2B,D), indicating that gaboxadol alone does not affect olfactory receptor neuron morphology. In axotomized flies, GFP signal was reduced in the antennal glomeruli, antennal lobe commissure, and olfactory receptor neuron axons at all time points (Fig 2C, Fig S2 E,I,M). In axotomized flies treated with gaboxadol, the GFP signal in the antennal glomeruli, antennal lobe commissure, and Or22a axons was significantly lower than in undisturbed axotomized flies (Fig 2E-H), and these findings were confirmed by monitoring the GFP signal over a time course spanning 3 days after ablation (Fig S2C–R). Taken together, these results reveal that pharmacological induction of sleep accelerates the engulfment of axotomized neurons by ensheathing glia.

**Figure 2.**
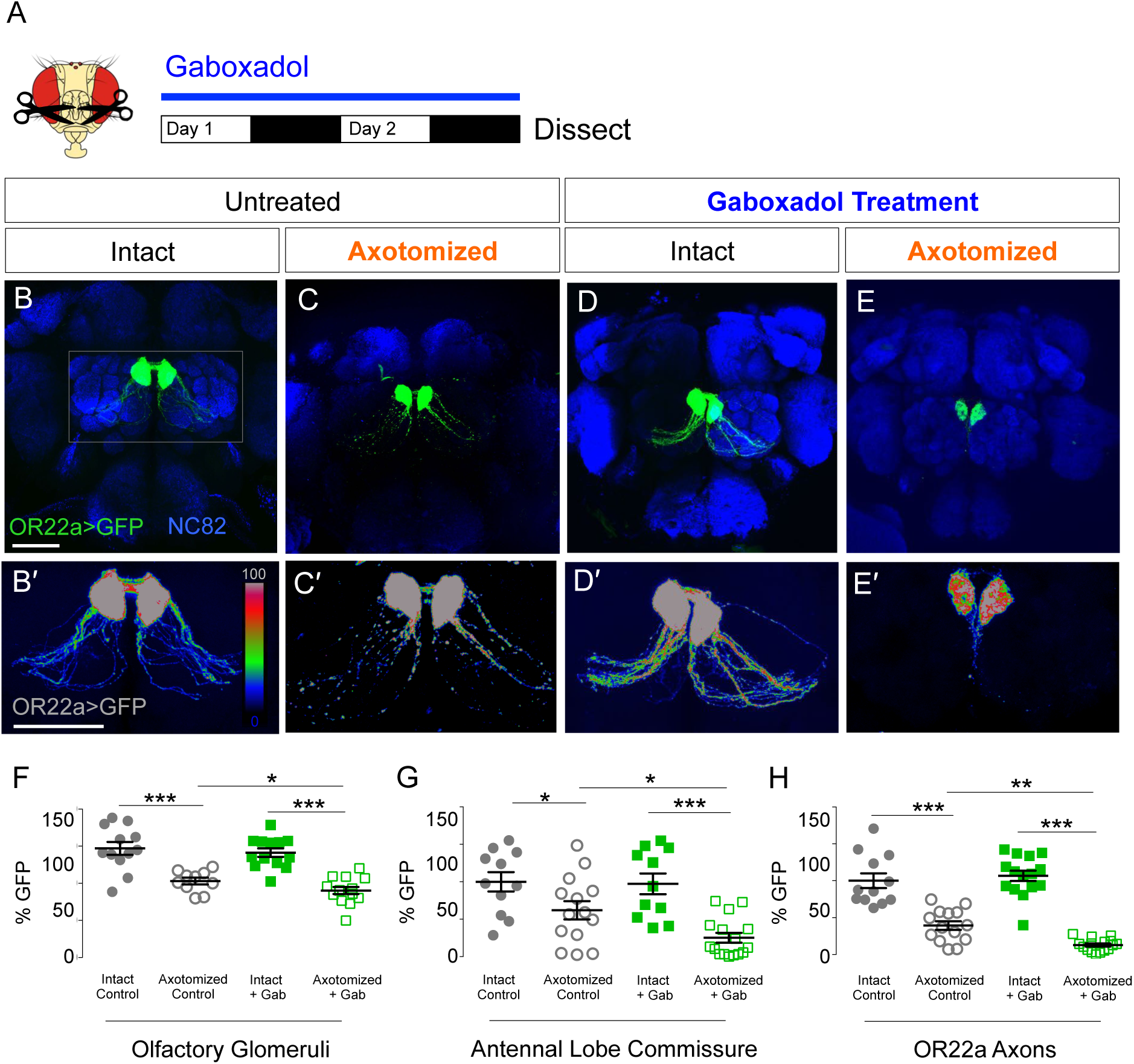
Sleep accelerates clearance of degenerating axons. **A)** Schematic of antennal ablation and sleep-promoting gaboxadol treatment. **(B–E)** Max intensity projection of GFP-labeled olfactory OR22a axons (green) with NC82 neuropil counterstain (blue). **(B)** OR22a neurons in unharmed control flies. **(C)** Flies with third antennal segment ablation and no drug treatment. **(D)** Gaboxadol sleep-promoting treatment in unharmed flies. **(E)** Flies with ablation plus gaboxadol. **(B′–E′)** Zoom and max intensity rainbow plots. **(F–H)** Both axotomized controls and flies subjected to ablation plus gaboxadol treatment exhibited a reduction in the GFP signal relative to unharmed individuals, but axotomized flies treated with gaboxadol exhibited faster removal of GFP than axotomized controls in OR22a glomeruli **(F**; ANOVA, F_(3,45)_=21.40, P<0.0001), antennal lobe commissure (**G**; ANOVA, F_(3,45)_=10.54, P<0.0001) and the axons (**H**; ANOVA, F_(3,45)_=53.59, P<0.0001). Tukey’s multiple comparisons tests: *P<0.05, **P<0.01, ***P<0.001. Scale Bar = 50 μm.

Axotomy of olfactory receptor neurons activates ensheathing glia, resulting in their innervation of the antennal lobe. To directly measure glial innervations following axotomy, we labeled glial membranes by genetic expression of membrane-bound GFP (mD8:GFP) under the control of the pan-glial driver Repo [19], allowing quantification of florescence within the antennal lobes (Fig 3A). Consistent with previous findings, antennal axotomy significantly increased membrane innervation of the antennal lobes (Fig 3B–D). In sleep-deprived flies, glial innervation of the antennal lobe was reduced following axotomy compared to undisturbed axotomized flies (Fig 3E,F, Fig S3A). Conversely, 2 days after axotomy, glial innervation of the antennal lobe was greater in gaboxadol-treated flies than in undisturbed controls (Fig 3G-L, Fig S3B). Therefore, sleep directly regulates membrane innervation of antennal glomeruli following injury.

**Figure 3.**
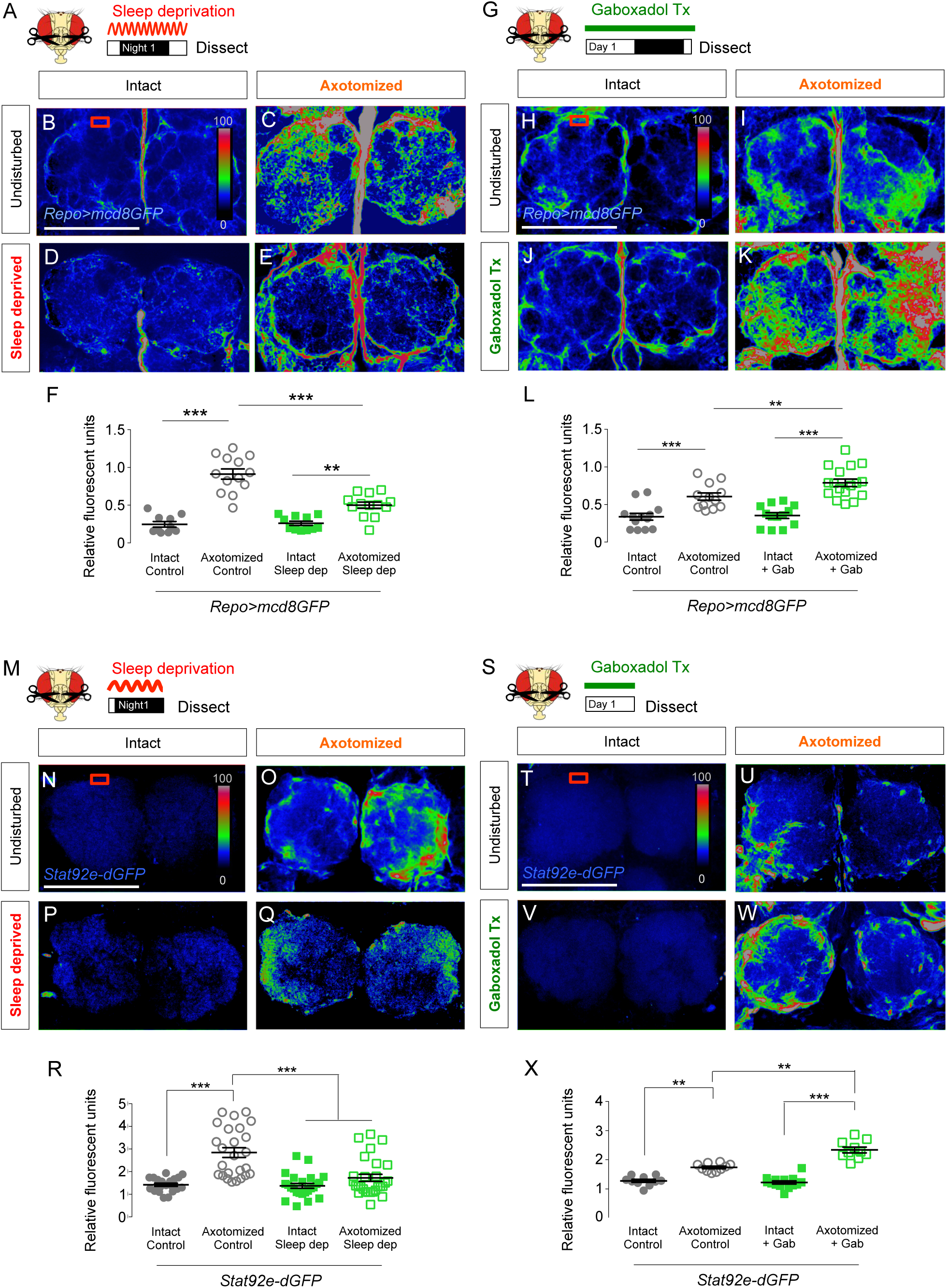
Glial membrane response in sleep-deprived and gaboxadol-treated flies. **(A)** Schematic of antennal ablation and sleep deprivation treatment for the GFP-labeled glial membrane experiments. **(B–E)** Rainbow intensity plots of glial membranes in intact individuals, either undisturbed (**B**) or sleep-deprived (**D**), and axotomized flies either undisturbed (**C**) or sleep-deprived (**E**). Red box indicates the glial membrane region that was quantified. Axotomized controls exhibited an increase in Repo-labeled glial membranes following antennal ablation (**C**). Axotomized flies that were sleep-deprived exhibited a reduced glial membrane response (**E**). Quantification of Repo-labeled glial membrane (**F**; ANOVA, F_(3,43)_=42.20, P<0.0001). **(G)** Schematic of antennal ablation and sleep-promoting gaboxadol treatment for the GFP-labeled glial membrane experiments. **(H–K)** Rainbow intensity plots of glial membranes in intact individuals without drug (**H**) or treated with gaboxadol (**I**), and axotomized flies without drug (**J**) or treated with gaboxadol (**K**). Axotomized controls exhibited an increase in Repo-labeled glial membranes after antennal ablation (**I**). Axotomized flies treated with gaboxadol exhibited a greater glial membrane response (**K**). Quantification of glial membranes (**L**; ANOVA, F_(3,41)_=23.08, P<0.0001). **(M)** Schematic of antennal ablation and sleep deprivation treatment for the *Stat92e-dGFP* experiments. **(N–Q)** Rainbow intensity plots of glial membranes in intact individuals either undisturbed (**N**) or sleep-deprived (**P**), and axotomized flies either undisturbed (**O**) or sleep-deprived (**Q**). **(R)** Axotomized controls exhibited an increase in the *Stat92e-dGFP* response following antennal ablation (**C**). Axotomized flies that were sleep-deprived exhibited a delayed response (ANOVA, F_(3,93)_=19.29, P<0.0001). **(S)** Schematic of antennal ablation and gaboxadol treatment for the *Stat92e-dGFP* experiments. **(T–W)** Rainbow intensity plots of glial membranes in intact individuals without drug (**T**) or treated with gaboxadol (**V**), and axotomy flies without drug (**U**) or treated with gaboxadol (**W**). **(X)** Axotomy flies exhibited a higher *Stat92e-dGFP* response, and this effect was stronger when they were treated with gaboxadol (ANOVA, F_(3,40)_=64.49, P<0.0001). Tukey’s multiple comparisons tests: *P<0.05, **P<0.01, ***P<0.001. Scale Bar = 50 μm.

To complement our analysis of glial membranes, we quantified signaling associated with the glial response to axonal injury. Activation of glia by axonal injury is marked by an upregulation of the transcription factor Stat92E [20]. The Stat92E::dGFP reporter mimics the endogenous expression pattern, providing a readout of injury-induced activity [20,21]. Due to the robust induction of Stat92E::dGFP following injury, and a ceiling effect at later time points, we measured Stat92E::dGFP signal flies 12 hrs after ablation. Reporter levels were significantly lower in flies sleep-deprived throughout the night than in undisturbed axotomized controls (Fig 4M-R, Fig S3C). Conversely, inducing sleep with gaboxadol for the 12-hour day following axotomy significantly increased Stat92E::dGFP signal relative to that in axotomized flies fed solvent alone(Fig 4S–X, Fig S3D). Taken together, these findings support the idea that sleep promotes the glial response to axonal injury.

**Figure 4.**
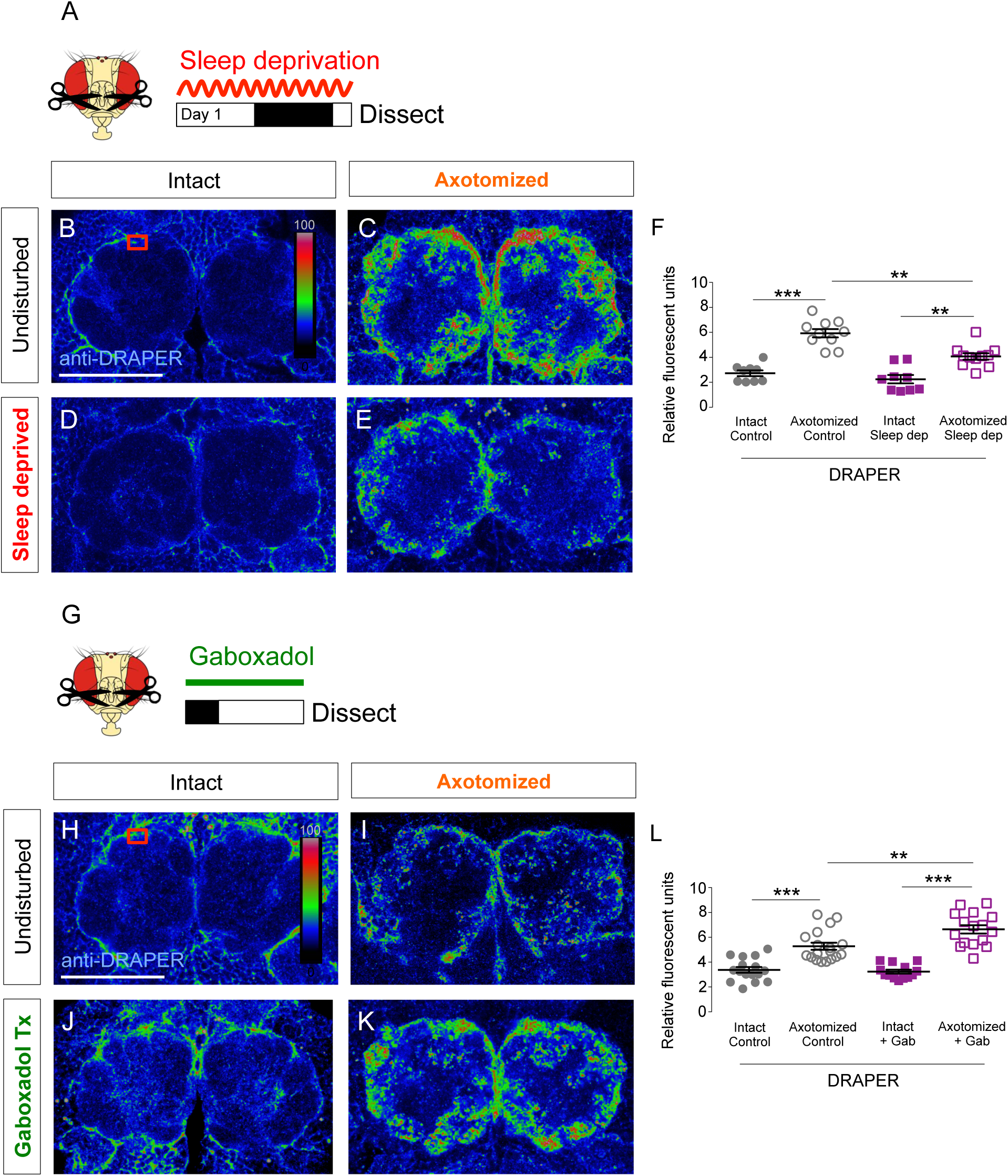
Draper response is affected by sleep state. **(A)** Schematic of the antennal ablation and sleep deprivation paradigm for th Draper experiments. **(B–E)** Rainbow intensity plots of anti-DRAPER in intact individuals either undisturbed (**B**) or sleep-deprived (**D**), and axotomized flies either undisturbed (**C**) or sleep-deprived (**E**). Axotomized controls exhibited a dramatic increase in DRAPER expression following antennal ablation (**C**). Although axotomized and sleep-deprived flies exhibited an increase in Draper levels, the response was reduced (**E**). Quantification of DRAPER expression (**F**; ANOVA, F_(3,35)_=30.98, P<0.0001). **(G)** Schematic of antennal ablation and gaboxadol treatment for DRAPER experiments. **(H–K)** Rainbow intensity plots of anti-DRAPER in intact flies without drug (**H**) or treated with gaboxadol (**J**), and axotomized flies without drug (**I**) or treated with gaboxadol (**K**). Axotomized controls show a dramatic increase in DRAPER expression following antennal ablation (**I**). However, axotomizeed flies treated with sleep-promoting gaboxadol exhibited an greater DRAPER response (**K**). Quantification of DRAPER expression (**L**; ANOVA, F_(3,52)_=34.95, P<0.0001). Tukey’s multiple comparisons tests: **P<0.01, ***P<0.001

Wallerian degeneration induces expression of the engulfment receptor Draper within ensheathing glia, which is essential for engulfment of severed axons [12,15]. To determine whether sleep regulates Draper function, we measured the effects of a loss or gain of sleep on Draper level within the olfactory ensheathing glia. To specifically measure the effects of sleep loss, we mechanically sleep-deprived flies for 3 days following ablation, and then immunostained for Draper (Fig 4A). As previously described, ablation resulted in robust Draper signal near and within the antennal lobe (Fig 4B–C) [12]. Sleep deprivation had no effect on Draper signal in intact flies (Fig 4A-D), but Draper levels within the antennal lobe were lower in axotomized flies than in undisturbed flies, suggesting sleep deprivation inhibits Draper-mediated engulfment (Fig 4D–F). To determine whether sleep promotes Draper localization within the antennal lobes, we treated flies with gaboxadol for 2 days following ablation, and then measured Draper levels. Gaboxadol treatment did not affect Draper levels in intact controls, but significantly increased Draper levels in axotomized flies relative to undisturbed flies (Fig 4H–L). Therefore, sleep promotes Draper localization to the glial innervations of the antennal lobe following axotomy.

Our findings reveal that sleep regulates Wallerian degeneration of fruit fly olfactory receptor neurons. We demonstrated that sleep promotes expression of multiple markers of glial activation, including membrane innervation antennal lobe and upregulation of Draper and Stat92E. Conversely, these markers were inhibited in sleep-deprived flies. In mammals, microglia perform many of the engulfing functions that are attributed to ensheathing glia in flies [6,13], and multiple lines of evidence suggest that microglia are regulated by sleep state. Microglia-mediated clearance may be critical for maintaining brain function: activation of microglia limits progression of Alzheimer’s disease (AD) in mouse models, suggesting that clearance of damaged neurites may be critical for health [25]. The functional link between sleep and glial function in fruit flies provides a platform for investigating the molecular basis for interactions between sleep and glia-mediated engulfment.

Although this study focused on how sleep affects the functions of olfactory ensheathing glia, it is also possible that sleep state impacts the damaged neurites themselves, in turn influencing the rate of degeneration. Numerous factors within injured neurons promote engulfment: for example, the biosynthetic enzyme nicotinamide mononucleotide adenylyltransferase (Nmat1) and its redox metabolite NAD+ play a critical prevent Wallerian degeneration [12,26]. Following axotomy, depletion of NAD+ results in activation of the *Drosophila* Toll receptor adaptor Sarm1, which in turn promotes engulfment [27]. Genetic approaches in the fly have identified numerous additional neural factors that modulate engulfment, including the zinc finger transcription factor Pebbled and the E3 Ubiquitin ligase Highwire, which promote degeneration [27–29], whereas the MAP Kinase Wallenda protects damaged axons from engulfment [30].

The observation that sleep promotes engulfment of damaged neurites reveals an additional role for sleep in the regulation of behavior and brain physiology. In flies, sleep is regulated by life-history traits including age, diet, and social interactions [31–34], Sleep is disrupted in numerous models of neurodegenerative disease, and promotes survival in response to oxidative stress [35–37]. Further, genetic or pharmacological induction of sleep alleviates disease progression in fly models of Alzheimer’s Disease, suggesting that sleep is neuroprotective [16,38,39]. Both sleep and Wallerian degeneration are disrupted in old flies, raising the possibility that restoration of sleep in old animals could protect against age-related decline in glial function [8,36,40]. The identification of sleep-dependent changes in the function and morphology of ensheathing glia will facilitate studies of the effects of sleep on glial function in aging and disease models.

## Materials and Methods

### Fly stocks and maintenance

Flies were reared and maintained on a 12:12 light–dark cycle in humidified incubators at 25°C and 65% humidity (Percival Scientific, Perry, IA, USA). The following fly lines were obtained from the Bloomington (BL) Stock Center: UAS-*mcD8GFP* (P{10XUAS-IVS-mCD8::GFP}; BL#32186 [41]; *Repo*-GAL4 (P{GAL4}repo/TM3, Sb1}; BL#7415; [19]), *Stat92e-dGFP* (P{10XStat92E-DGFP}; BL#26199; [21]), OR22a::GFP (Or22a-Mmus\Cd8a.GFP}; BL#52620 [42]. The background control line used in this study was *w*^*1118*^ (BL# 5905; [43]) and all experimental stocks were either generated in this background or outcrossed to *w*^*1118*^ for at least six generations prior to analysis. Mated females were aged for 2–4 days in standard vials prior to the experiments.

### Sleep deprivation

Axotomized and intact control flies were sleep-deprived in groups of 8–10 flies in housing vials mounted on a vortexer (Scientific Industries, Inc, Vortex Genie 2, #SI-0236). Mechanical shaking stimulus was applied using a repeat-cycle relay switch (Macromatic, #TR631122); flies were shaken for ∼3–4 seconds every minute for the duration of the experiment, as previously described [44,45]. Sleep deprivation began following axotomy and recovery at ∼ZT8. During sleep deprivation, flies were maintained on a 12:12 hr light:dark cycle at 25°C. Undisturbed control flies were maintained under identical conditions in the absence of vortexing.

### Gaboxadol treatment

Gaboxadol (4,5,6,7-retrahydroisoxazolo[5,4-*c*]pyridin-3-ol hydrochloride, THIP hydrochloride; Sigma Aldrich, St. Louis, MO, USA, # T101) was dissolved in dH_2_O at 1 mg/ml, then mixed to 0.01 mg/ml in cooled, standard fly food (Bloomington Formulation, Nutri-fly, #66-113, Genesee Scientific, San Diego, CA, USA), as previously described [16,46]. Axotomized and intact control flies were placed on gaboxadol-laced food immediately following axotomy, where they remained on a 12hr light/12 hr dark cycle at 25°C for the duration of the experiment.

### Immunohistochemistry and imaging

Brains were dissected in cold phosphate buffer solution (PBS) and fixed in 4% paraformaldehyde in PBS containing 0.5% Triton-X100 [PBT], for 40 minutes at room temperature (RT). After three washes (5 min each) with PBT and rinsing overnight at 4°C, the brains were incubated with relevant primary antibody, anti-BRP (1:5; #nc82, AB_2314866, Developmental Studies Hybridoma Bank) or anti-DRAPER 5D14 (1/500; #draper, AB_2618105, Developmental Studies Hybridoma Bank), for 48 hr at 4°C with rotation. After 3 10-min washes in PBT at RT, the brains were incubated with secondary antibody for 90 min at RT: (Alexa Fluor 555 donkey anti–mouse IgG, Invitrogen, Carlsbad, CA, USA, #A31570), and then washed three times (30 min each) in PBT, followed by a final overnight wash in PBT. Brains were mounted in Vectashield (H1000, Vector Labs, Burlingame, CA, USA). Fluorescence images were acquired on a Nikon A1 laser-scanning confocal microscope. Brains were imaged using the 561 nm and 488 nm lasers sequentially, with imaging settings (e.g., laser power, gain, and zoom) kept constant throughout the entire experiment. Image stacks were acquired using the ND Sequence Acquisition function, imaging from the above to below the antennal lobes using a 2-μm step size.

### Quantification and statistical analyses

Quantification of *OR22a>mcD8GFP* (Figures 1 and 2) was performed by generating a maximum intensity projection of the OR22a-label neurons and quantifying the sum GFP intensity in each of the three regions of interest (ROI): antennal lobe commissure (ALC), glomeruli, and axons using the Nikon AR Image Analysis software (Nikon, Melville, NY, USA; see ROIs in Figure S1C), as previously described [12]. *Repo>mcD8GFP, State92e-dGFP*, and anti-DRAPER (Figures 3 and 4) were quantified using two approaches. The first quantification method using a standardized ROI of the outer antennal lobe region (minus background measurement) from a single z-slice, was performed using FIJI as previously described [12,47]. A second measurement was performed by generating a sum of maximum intensities from the antennal lobe stack, and then quantifying the integrated density measurement of entire antennal lobe as an ROI, again using FIJI. Statistical comparisons were determined by one-way or two-way ANOVA with Tukey’s correction for multiple comparisons, using the InStat software (GraphPad Software v.6.0). The two-tailed P-value for significance was P<0.05. “n” refers to the number of biological samples tested, unless stated otherwise.

### Fly Sleep Behavior

Sleep behavior was measured using the *Drosophila* Activity Monitoring system (DAMs; DAM2 system, Trikinetics), which detects activity by monitoring infrared beam crossings of single flies. The number of beam-breaks is used to determine the amount of sleep, along with associated sleep metrics, by identifying 5-min bouts of quiescence using the *Drosophila* Sleep Counting Macro [48–50]. Flies were briefly anesthetized with CO_2_ and loaded individually in DAMs tubes with standard fly food (Bloomington Formulation, Nutri-fly, #66-113, Genesee Scientific). Sleep was then measured for 12 hr (either 12-hr light or 12-hr dark), depending on the relevant comparison. During the experiment, flies were maintained in incubators (Model #DR-41NL, Percival) at 25°C with ∼50% relative humidity, in accordance with previously published methods. All behavioral experiments utilized 4–7-day old mated female flies.

### Three-dimensional reconstruction

Confocal stacks were processed through FIJI ImageJ v1.51s, and then converted to NIFTI files for analysis using the Amira software, v6.4 (FEI) as previously described [1]. Files were opened in Amira and visualized using the Volren 3D render module. All visualization histograms were equalized to 110–4095 to ensure consistency across samples. All Volren modules were co-animated with the Animate Ports module attached to each image stack, so that all three-dimensional rotations and scaling were the same in all samples. Videos were rendered by attaching a camera orbit to the Volren renderings in the Animation editor. Time values were set to rotate around the z-axis 360° from beginning to end. Files were saved as 720p .avi videos.

## Acknowledgements

The authors are grateful to Mary Logan (OHSU) for guidance and discussion, and Melanie Gratton (FAU) for technical assistance. This work was supported by National Institutes for Health (NIH) Grants R01NS085252 and R01HL143790 to ACK.

## Author contributions

The experiments were designed by BAS and ACK. All authors contributed to writing the manuscript. Experiments were performed by BAS and JBJ.

## Supplemental figure legends

**Figure S1.**
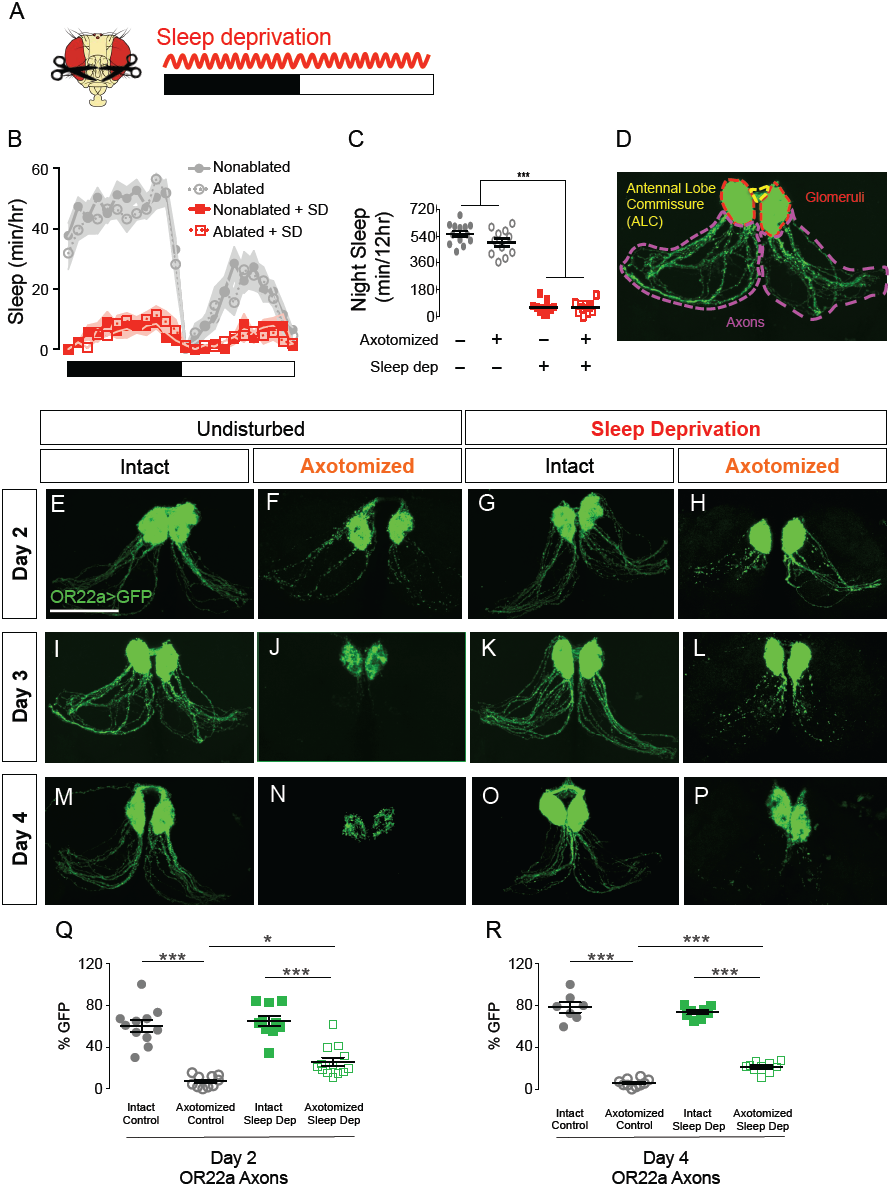
Sleep deprivation slows the rate of Wallerian degeneration. **(A)** Schematic of antennal ablation and sleep deprivation treatment. (**B)** Sleep profile of intact (solid line) or axotomized (dashed line) flies either undisturbed (grey) or sleep-deprived (red). **(C)**. Control flies with antennal ablation slept normally, and axotomized sleep-deprived flies responded to mechanical sleep deprivation as anticipated (ANOVA, F_(3,40)_=187.2, P<0.0001). **(D)** Regions of interest (ROIs) of OR22a neurons were quantified. **(E–P)** Time series showing degeneration under different treatments: 2 days post-ablation (**E–H**), 3-days post ablation (**I–L**), and 4 days post-ablation (**M–P**). Quantifications are shown in **Q–R (**Day 2: ANOVA, F_(3,40)_=38.89, P<0.0001; Day 2: ANOVA, F_(3,31)_=222.2 P<0.0001). Tukey’s multiple comparisons tests: *P<0.05, **P<0.01, ***P<0.001. Scale Bar = 50 μm.

**Figure S2.**
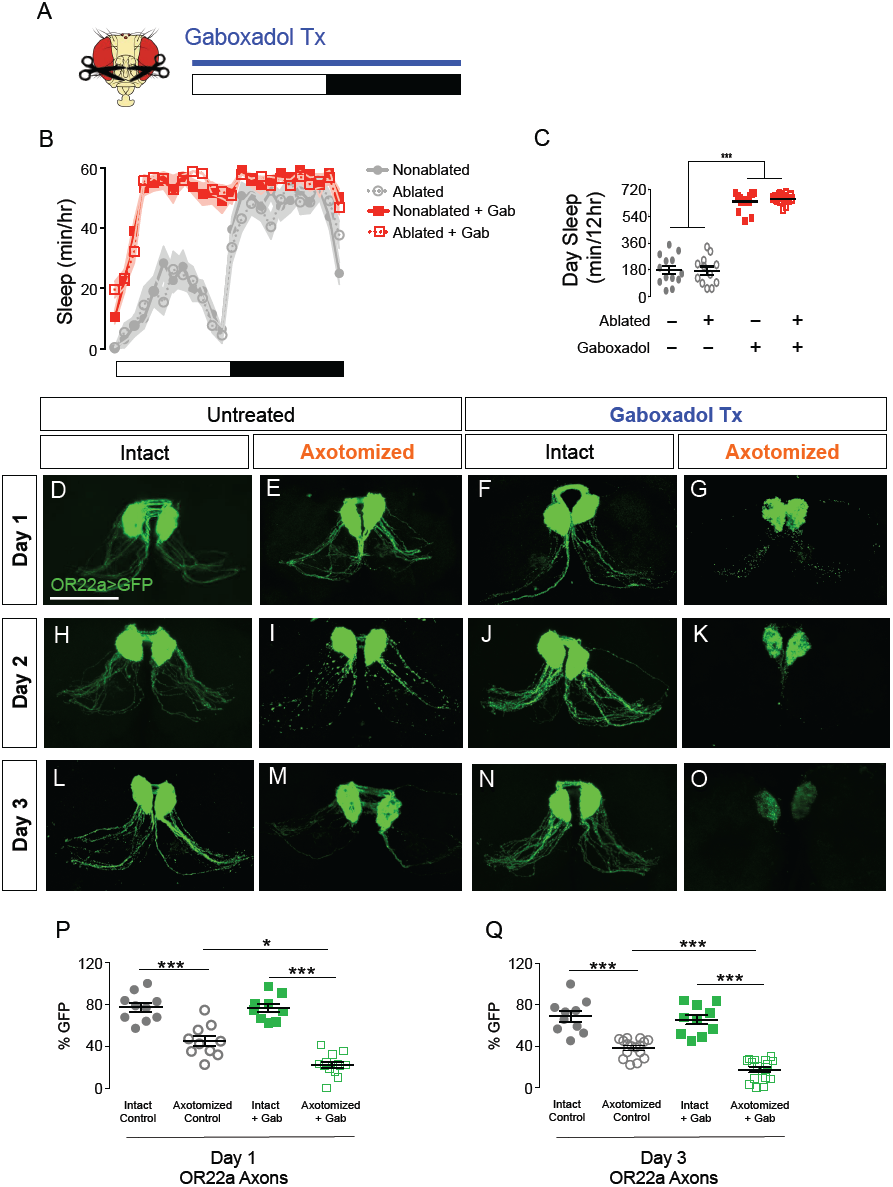
Sleep increases the rate of Wallerian degeneration. **A)** Schematic of antennal ablation and sleep-promoting gaboxadol treatment. (**B)** Sleep profile of intact (solid line) or axotomized (dashed line) flies either without drug (grey) or treated with gaboxadol (red). **(C)** Control flies with antennal ablation slept normally, and axotomized gaboxadol-treated flies increase sleep as anticipated (ANOVA, F_(3,46)_=162.4, P<0.0001). **(D–O)** Time series showing degeneration under different treatments: 1 day post-ablation (**D–G**), 2 days post-ablation (**H-K**), and 3 days post-ablation (**L-O**). Quantifications are shown in **P–Q** (**Day1:** ANOVA, F_(3,38)_=48.39, P<0.0001; Day 3: ANOVA, F_(3,48)_=51.05, P<0.0001). Tukey’s multiple comparisons tests: *P<0.05, **P<0.01, ***P<0.001. Scale Bar = 50 μm.

**Figure S3.**
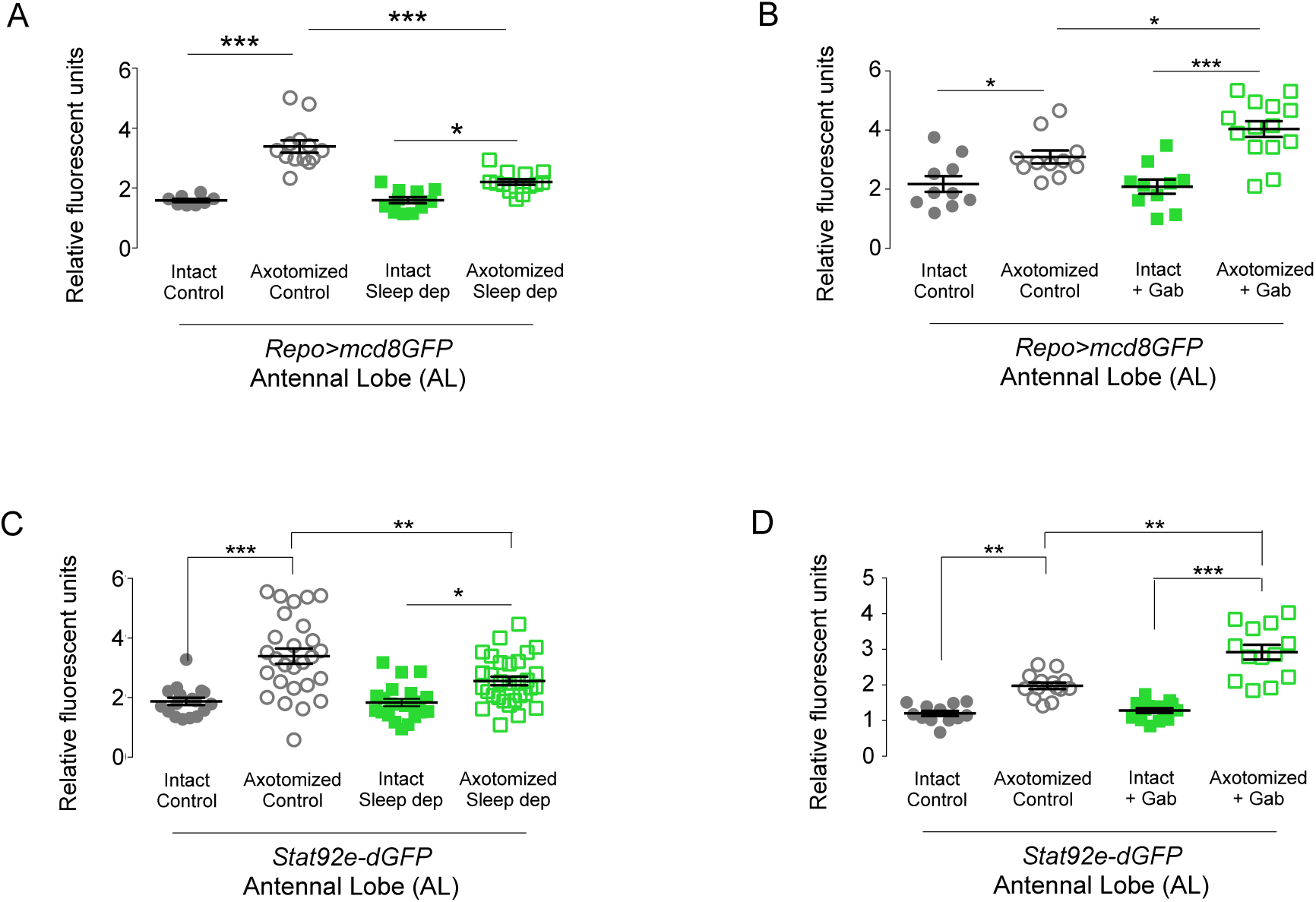
Glial membrane response in sleep-deprived and gaboxadol-treated flies. **(A–D)** Second method of quantification of GFP-labeled glial membranes and th *Stat92e-dGFP* response involved generating a sum intensity projection from the z-stack range that encompassed the antennal lobes, and quantifying the integrated density measurement from the entire area of the antennal lobe (AL). **(A–B)** Repo-labeled glial membrane quantification of the entire AL under sleep deprivation (**A**; ANOVA, F_(3,43)_=36.27, P<0.0001), and gaboxadol treatment (**B**; ANOVA, F_(3,41)_=13.81, P<0.0001). **(C-D)** *Stat92e-dGFP* quantification of the entire AL under sleep deprivation (**C**; ANOVA, F_(3,93)_=15.54, P<0.0001), and gaboxadol treatment (**D**; ANOVA, F_(3,40)_=41.56, P<0.0001). Tukey’s multiple comparisons tests: *P<0.05, **P<0.01, ***P<0.001. Scale Bar = 50 μm.

**Figure S4.**
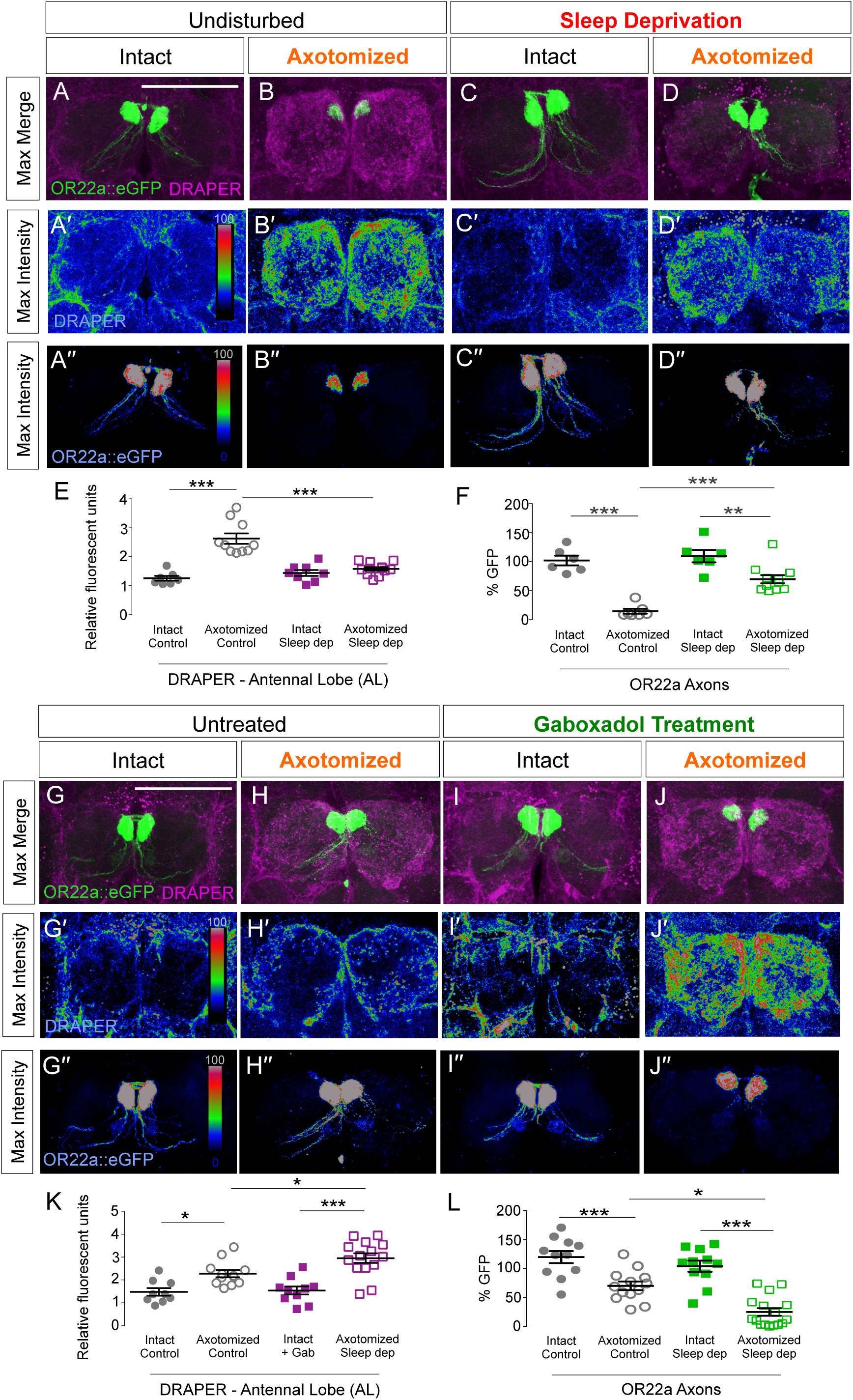
Sleep deprivation impedes the Draper response. (**A-E**) Maximum intensity projection, merged image of GFP-labeled OR22a neurons (green) counterstained with anti-DRAPER (magenta) in intact individuals either undisturbed (**A**) or sleep-deprived (**C**), and axotomized flies either undisturbed (**B**) or sleep-deprived (**D**). **(A′–D′)** Rainbow intensity plots of anti-DRAPER. **(A′′-D′′)** Rainbow intensity plots of GFP-labeled OR22a neurons. **(E)** Second quantification of anti-DRAPER for entire antennal lobe (AL; ANOVA, F_(3,35)_=26.64, P<0.0001).**(F)** Quantification of GFP-labeled OR22a axons in brains co-stained with anti-DRAPER (ANOVA, F_(3,26)_=27.21, P<0.0001). (**G–J**) Maximum intensity projection, merged image of GFP-labeled OR22a neurons (green) counterstained with anti-DRAPER (magenta) in intact individuals without drug (**G**) or treated with gaboxadol (**I**), and axotomized flies without drug (**H**) or treated with gaboxadol (**J**). **(G′–J′)** Rainbow intensity plots of anti-DRAPER. **(G′′–J′′)** Rainbow intensity plots of GFP-labeled OR22a neurons. **(K)** Second quantification of anti-DRAPER for entire antennal lobe (AL; ANOVA, F_(3,52)_=14.62, P<0.0001). **(L)** Quantification of GFP-labeled OR22a axons in brains co-stained with anti-DRAPER (ANOVA, F_(3,47)_=27.63, P<0.0001). Tukey’s multiple comparisons tests: *P<0.05, **P<0.01, ***P<0.001. Scale Bar = 50 μm.

**Supplemental Video 1: 3D reconstruction of axotomized OR22a neurons with sleep deprivation**

